# Ultra-low-illumination, high-fidelity longitudinal monitoring of cerebral perfusion via deep learning–enhanced laser speckle contrast imaging

**DOI:** 10.64898/2026.03.10.710928

**Authors:** Mingliang Xu, Fangyuan Li, Guanyi Zhu, Haoran Ma, Fei He

**Author notes:** These authors contributed equally (Mingliang Xu, Fangyuan Li).

## Abstract

Laser Speckle Contrast Imaging (LSCI) is a non-contact, label-free optical technique widely used in biomedical research and clinical applications. It enables real-time visualization and quantification of microvascular blood flow by analyzing the temporal fluctuations of laser speckles induced by moving red blood cells. However, conventional LSCI uses visible or near-infrared illumination, which—during prolonged exposure (*e.g*., >1□hr)—can induce sublethal neural stress and cause signal drift, compromising physiological relevance and raising ethical concerns. To mitigate these limitations, we introduce TunLSCI—a TransUNet-based recovery network designed to reconstruct high-fidelity mouse cerebral blood flow (CBF) indices from ultra-low-illumination LSCI. We train our network on paired ultra-low-illumination (1.27 µW/mm^2^) and conventional LSCI data (∼200 µW/mm^2^ illumination, the latter as reference), and demonstrate that it outperforms the conventional standard analytical LSCI processing pipeline based on stLASCA, particularly in reconstructing fine vasculature from few frames, suppressing speckle noise, and maintaining robustness against exposure variations. We validate that the proposed TunLSCI reduces illumination power density by ∼157-fold compared with conventional stLASCA, well below the safety threshold for cortical exposure in mice and markedly improves stability during a 2-hour continuous mouse CBF monitoring. Our method significantly minimizes the phototoxic burden of LSCI while preserving spatiotemporal fidelity and quantitative accuracy, thus enabling longitudinal, high-biosafety cerebral perfusion tracking *in vivo* over multi-hours.

## 1. Introduction

Laser speckle contrast imaging (LSCI) is a wide-field, label-free optical technique that provides real-time, 2D blood flow maps by analyzing laser speckle blurring caused by moving red blood cells [1, 2]. It computes spatial or temporal contrast—typically the ratio of pixel intensity standard deviation to mean—to generate high-resolution perfusion maps reflecting relative flow velocity and density [3-5]. Due to its simplicity, speed, and sensitivity, LSCI is well-validated in preclinical applications, including cerebral blood flow (CBF) monitoring, skin and retinal microcirculation assessment, and pharmacological studies. For instance, it captures rapid, stimulus-evoked perfusion changes during neurovascular coupling studies in awake or anesthetized animals [6]. Its portability, low cost, and compatibility with standard surgical microscopes support clinical translation — especially intraoperative CBF mapping during tumor resection or epilepsy surgery, where it delivers real-time feedback on tissue viability, vascular integrity, and intervention effects (*e*.*g*., temporary clipping or cortical stimulation) [7-9]. Optical strategies that extend the effective depth of field have also been introduced to improve *in vivo* LSCI under practical imaging geometries [10].

Conventional LSCI uses visible-to-near-infrared lasers (630–850 nm) for *in vivo* microvascular blood flow imaging, as this range optimizes tissue penetration and scattering for superficial applications. However, prolonged LSCI acquisition—common in longitudinal studies of neurovascular dynamics, ischemia-reperfusion, or drug interventions—requires sustained laser illumination, risking cumulative photochemical damage to cortical tissue. This damage arises from photon absorption by hemoglobin, melanin, and cytochrome c oxidase, leading to localized heating and ROS generation [11, 12]. It can also induce reversible but functionally relevant neural stress such as transient membrane potential shifts, calcium dysregulation, and mild mitochondrial dysfunction—disrupting baseline neurovascular coupling. These physiological deviations manifest as time-dependent signal drift in LSCI: speckle contrast decreases not only due to hemodynamics, but also from progressive microstructural alterations that alter light scattering. Such confounds compromise longitudinal fidelity and quantitative perfusion accuracy, especially in temporally sensitive experiments like sensory stimulation or pharmacological challenge.

Several strategies have been proposed to alleviate laser-induced physiological disturbances during prolonged LSCI. For instance, incorporating nicotinamide adenine dinucleotide (NADH) fluorescence imaging with LSCI can detect metabolic stress and early phototoxicity before morphological damage occurs. Recent work has also integrated NIR fluorescence with LSCI (with optical tissue clearing for transcranial readout) to co-monitor vascular morphology and flow under light–tissue interactions [13]. In contrast, metabolic-stress readout based on NAD(P)H autofluorescence typically relies on UV excitation (∼340– 390 nm), which requires specialized optics/sensors, limits penetration, and increases system complexity. Speckle entropy feedback algorithms dynamically adjust laser power by analyzing the spatiotemporal heterogeneity to identify early tissue stress before it becomes visible for unsupervised long-term studies, but demand high-speed processing (challenging for resource-limited systems) and tissue-specific entropy baselines requiring prior validation [14]. For clinical application, an intraoperative 5-minute on/2-minute off cycling protocol minimizes cumulative thermal load, facilitates tissue recovery, and preserves diagnostic utility without hardware changes. Yet, intermittent acquisition may miss rapid hemodynamic transitions, and dosing must be tailored to individual patient pathophysiology.

To overcome the above limitations, we depart from conventional high-illumination LSCI protocols and instead acquire LSCI data under ultra-low illumination—up to two orders of magnitude lower than standard settings—well below reported safety thresholds for mouse cortical exposure. While this substantially reduces photodamage risk and improves animal-welfare compliance, the reduced photon budget markedly degrades the speckle SNR, leading to noisy, low-contrast perfusion maps that are unreliable for quantitative perfusion indexing with standard analytical processing. To recover robust, high-fidelity CBF indices in this photon-limited regime, we adopt the TransUNet architecture as a learning-based recovery backbone and introduce a minimal input-adaptation layer to accommodate an n-frame ultra-low-illumination speckle stack concatenated along the channel dimension. Our TransUNet-based LSCI (TunLSCI) learns a nonlinear mapping from an n-frame ultra-low-illumination raw speckle stack concatenated along the channel dimension to reference blood-flow index (BFI) targets computed from paired standard-illumination recordings using conventional standard analytical LSCI processing pipeline spatiotemporal laser speckle contrast analysis (stLASCA). TunLSCI retains the original TransUNet encoder–decoder transformer backbone and only introduces a minimal input-adaptation layer to support the multi-frame input, thereby enabling direct utilization of short temporal context from a small number of frames, even in the single-frame setting. Quantitative evaluation demonstrates consistent improvements in PSNR, SSIM, and vessel-structure fidelity, supporting reliable microvascular perfusion assessment at illumination levels as low as 1–5% of standard acquisition. Our method significantly reduces the LSCI-associated phototoxicity, enhancing the stability and reliability of CBF measurements during extended, hours-long continuous monitoring in preclinical studies.

## 2. Methods

### 2.1 Subjects and animal surgery

A total of three female and two male C57BL/6J wild type mice were used in the experiments. The animals’ age was 21-24 weeks at the cranial window surgery. They were housed individually after surgery at a 12-hr, 7:00 to 19:00, light-to-dark cycle. All animals received standardized cage supplementation (cage enclosures, nesting material and objects to gnaw) with water/food ad libitum. All animals received one surgery during which a chronic cranial window was mounted on the skull.

Mice were anesthetized with medical O2 vaporized isoflurane (3%) in an induction chamber and then placed prone in a stereotaxic frame (Stoelting, IL, USA) via nose-cone inhalation of medical O2 vaporized isoflurane at 1.5–2% (RWD, SZ, China). Dexamethasone (2 mg/kg) were administrated subcutaneously to reduce pain and inflammation during the craniotomy and implantation procedure. Body temperature was maintained at 37 °C with a feedback heat pad (Stoelting, IL, USA). Heart rate and breathing rate were monitored by the surgeon. A circular portion of skull (3 mm in diameter) atop the motor and somatosensory cortex was removed with a dental drill (Stoelting, IL, USA) under constant phosphate buffered saline flushing. Dura mater was removed to facilitate a larger imaging depth in the following imaging sessions. A 3 mm round cover glass was placed over the exposed cortex. Gentle pressure was applied to the cover glass while the space between the coverslip and the remaining skull was filled with Kwik-sil adhesive (World Precision Instruments, FL, USA). A layer of cyanoacrylate was applied to cover the skull and the Kwik-sil. Then an initial layer of C&B-Metabond (SUN MEDICAL CO., LTD, Shiga, Japan) was applied over the cyanoacrylate. This process ensured a sterile, air-tight seal around the craniotomy and allowed for restoration of intracranial pressure. A second layer of Metabond was used to cement a customized titanium head-plate for later head-constrained measurements.

The LSCI experiments started no sooner than 4 weeks after surgery to allow sufficient recovery of the animals [15-17]. All experiments were approved by the Institutional Animal Care and Use Committee (IACUC) at the Shanghai Institute of Optics and Fine Mechanics.

### 2.2 LSCI instrumentation, acquisition and processing

LSCI of CBF was performed using a 785-nm laser diode (70 mW; OBIS LX, Coherent Inc.) to illuminate the craniotomy at an oblique angle to monitor the mouse cerebral blood flow. The backscattered laser light was relayed to a complementary metal-oxide semiconductor camera (STC-MBA1002POE, Sentech) with 2.5× magnification and acquired 3856 × 2764-pixel frames for a field of view of 2.5 mm by 1.8 mm. The frame rate was 7.32 frames per second (fps) with a 3-ms exposure time (**Fig. 1a**). For each cranial-window location, we acquired back-to-back paired recordings in the same field of view (FOV) to preserve pixel-level correspondence while minimizing physiological mismatch (**Fig. 1b**): a low-power acquisition at 1.27 μW/mm^2^ with long exposure (60 ms or 120 ms) and a high-power reference acquisition at 200 μW/mm^2^ with 3 ms exposure. Each recording contained 1000 frames, and the switching latency between power settings was ∼1 s. The held-out evaluation cohort consisted of 5 mice, each with three cranial windows (15 window locations total); at each location, three recordings were acquired spanning two powers and two low-power exposures (45 recordings total). For all sessions of LSCI imaging, mice were awake and head-fixed on a customized, low-profile treadmill.

**Fig. 1.**
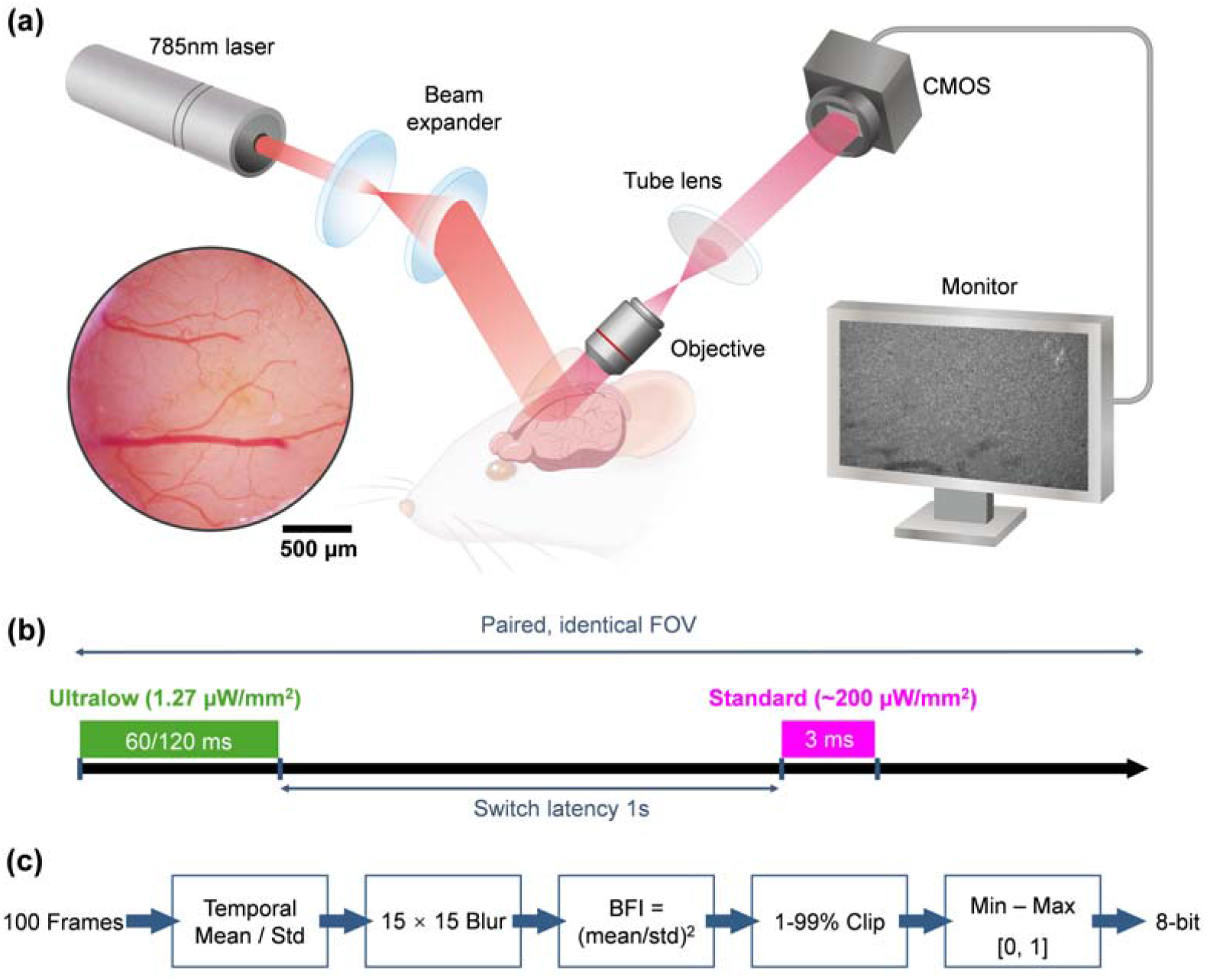
Paired low-exposure LSCI acquisition and learning-based recovery workflow. (**a**) Schematic of the home-built LSCI setup. Inset shows a photograph of the mouse cranial window 6 weeks after surgery. (**b**) Back-to-back paired recordings in the same FOV: low exposure (70 μW; 1.27 μW/mm^2^; 60/120 ms exposure time; 1000 frames) and standard-exposure reference (11 mW; 200 μW/mm^2^; 3 ms exposure time; 1000 frames); switching latency ∼1 s. (**c**) Reference BFI generation (stLASCA): 100-frame temporal statistics with 15×15 spatial smoothing; *BFI* = (µ/σ)^2^; 1–99% clipping; min–max normalization to [0,1]; 8-bit quantization.

Reference targets were generated from the conventional stLASCA under standard exposure (**Fig. 1c**) [18]. For a 100-frame window, we computed the per-pixel temporal mean and standard deviation σ, applied identical spatial smoothing to both moment maps using a 15 × 15 kernel, and formed an inverse-contrast-derived blood flow index surrogate as *BFI* = (*µ*/*σ*)^2^ (equivalently 1/*K*^2^ with speckle contrast *K* = *σ*/*µ*). The BFI map was then clipped to the 1st–99th percentiles, min–max normalized to [0,1], and quantized to 8-bit for training and evaluation. To restrict quantitative reporting to vasculature, vessel regions were manually delineated to create a vessel mask used for metric computation (and for consistent ROI definition across figures).

Reference BFI targets were generated using stLASCA under standard illumination (power density ≈ 200 μW/mm^2^, exposure time = 3 ms) (**Fig. 1c**). For a 100-frame temporal window, we computed per-pixel temporal mean (*µ*) and standard deviation (*σ*), applied spatial smoothing to both moment maps using a 15 × 15 kernel, and formed a BFI surrogate as (equivalently 1/*K*^2^ with speckle contrast *K* = *σ*/*µ*). The resulting BFI map was clipped at the 1st–99th percentiles, min–max normalized to [0, 1], and quantized to 8-bit for training and evaluation. To focus evaluation on vasculature, vessel regions were delineated by manual annotation to form a vessel mask used for metric computation and for consistent reporting across figures.

### 2.3 Neural network design and evaluation

For learning-based recovery (**Fig. 2**), the network input was a contiguous n-frame segment from the low-exposure speckle sequence (power density = 1.27 µW/mm^2^, exposure time = 60/120 ms), stacked along the channel dimension with *n* ∈{1, …,16}; the network output was a normalized [0,1] BFI map. Specifically, let *I*_*t*_ ∈ ℝ^*H*×*W*^denote the raw speckle intensity frame at time t. The network input was constructed by channel-wise concatenation of *n* consecutive frames,

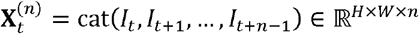

and the model prediction was 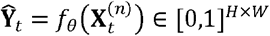, where *f*_*θ*_ (·)is the TransUNet-based recovery network parameterized by *θ*. We adopted a TransUNet-based architecture, where convolutional encoding captures multi-scale spatial features, patch embedding/tokenization enables transformer-based nonlocal modeling, and a U-shaped decoder reconstructs full-resolution perfusion maps [19]. Supervision used the aligned standard-illumination (*e*.*g*., power density ∼200 µW/mm^2^, exposure time = 3 ms) reference BFI **Y**_*t*_ ∈ [0,1]^*H*×*W*^ defined above. Training samples were generated by randomly sampling contiguous n-frame segments from each low-exposure recording while keeping paired low/high recordings grouped to avoid cross-split leakage. We trained separate models for each *n* to isolate the frame-budget effect. Optimization used a mean-squared-error objective between predicted and reference BFI,

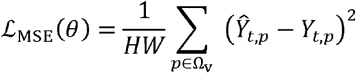

where Ω_v_ denotes the manually annotated vessel mask field and *p* indexes spatial locations. Validation performance was monitored to select the best checkpoint. For exposure-robustness evaluation (**Fig. 6**), models trained using 60-ms low-exposure inputs were tested on both 60-ms and 120-ms low-exposure recordings, while keeping the reference definition fixed.

**Fig. 2.**
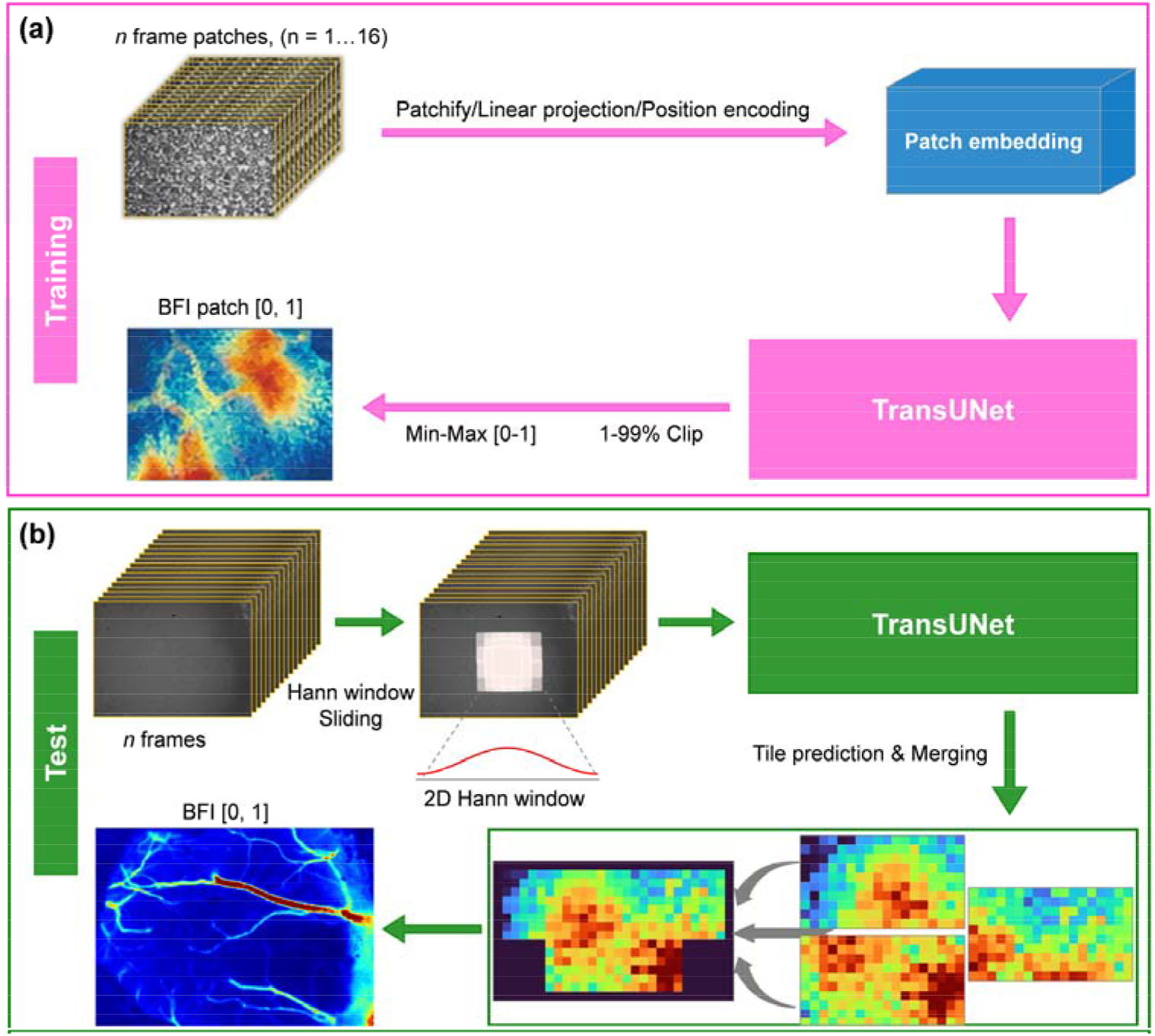
TransUNet-based LSCI (TunLSCI) training and Hann-window sliding inference. (**a**) Training pipeline: n-frame input stack (*n* = 1-16) → patch embedding/TransUNet → BFI output in [0,1]. (**b**) Testing pipeline: n-frame input stack → Hann-window sliding inference → tile prediction & Hann-weighted merging → full-field BFI output in [0,1].

Performance was quantified using PSNR, SSIM, and multiscale SSIM (MS-SSIM) computed on 8-bit BFI maps within the vessel mask [20, 21]. Both prediction and reference BFIs were quantized to 8-bit as 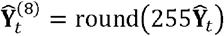 and 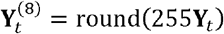. The vessel-masked mean squared error was computed as

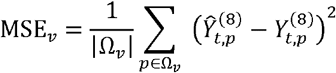

and PSNR was defined by

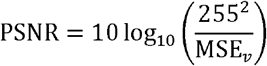

SSIM was computed between 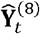 and 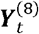 as

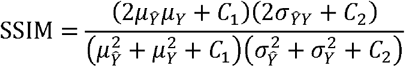

where 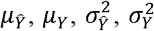, and σ_*ŶY*_ denote the mean, variance, and cross-covariance computed over vessel pixels, and *C*_1_, *C*_2_ are stabilizing constants. MS-SSIM was computed as a multi-scale product of luminance, contrast, and structure components. Photodamage-related perturbations (Fig. 3) were assessed by applying 2 h of standard illumination and comparing perfusion readouts to low-exposure measurements acquired in a separate recording session, with identical processing and visualization settings to ensure that observed differences reflect exposure-associated shifts rather than analysis inconsistencies.

**Fig. 3.**
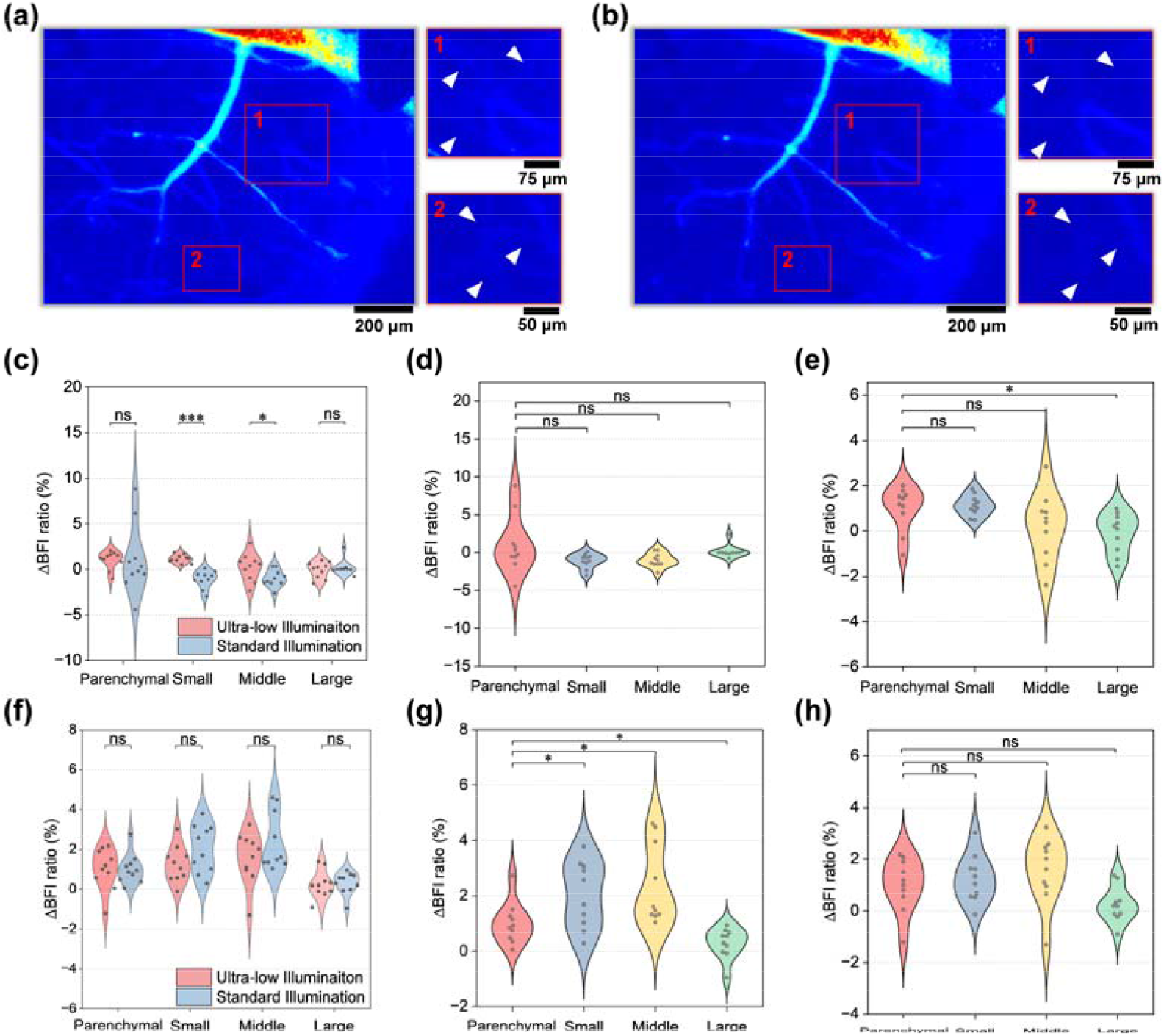
Prolonged standard illumination degrades microvascular perfusion readouts in LSCI. (**a, b**) Representative blood flow index (BFI) maps acquired under ultra-low illumination (a) and after prolonged standard illumination (b). Red boxes indicate regions of interest (ROIs), and the magnified views highlight distal microvessels (arrowheads). Compared with ultra-low illumination, prolonged standard illumination reduces the visibility of fine vascular branches, particularly in small-caliber vessels. (c-e) Venous ROI analysis. (**c**) Parenchyma-normalized ΔBFI distributions comparing ultra-low and standard illumination across parenchymal control, small (<10 μm), middle (10-30 μm), and large (>30 μm) vessel classes. (**d, e**) Class-wise ΔBFI distributions for venous ROIs under standard illumination (d) and ultra-low illumination (e), respectively. (f-h) Arterial ROI analysis. (**f**) Parenchyma-normalized ΔBFI distributions comparing ultra-low and standard illumination across parenchymal control, small (<10 μm), middle (10-30 μm), and large (>30 μm) vessel classes. (**g, h**) Class-wise ΔBFI distributions for arterial ROIs under standard illumination (g) and ultra-low illumination (h), respectively. Dots denote individual ROIs, violin plots show the data distributions, and statistical annotations are indicated in the plots.

### 2.4 CBF quantification and statistics

Because LSCI provides a relative perfusion surrogate (BFI) rather than absolute flow speed/flux/hematocrit, statistical analyses were performed on BFI-based CBF indices [22]. For ROI-based quantification, the mean BFI within each ROI was computed, and a parenchyma-normalized relative index was defined as 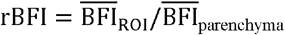. ROIs were stratified by tissue class (parenchyma vs vessel) and vessel type (artery vs vein). Vessel caliber was further grouped into three categories based on the measured vessel diameter on the reference map (e.g., small: <10 µm; middle: 10-30 µm; large: >30 µm). The parenchyma ROI was defined as the complement of the vessel mask within the field of view. For **Fig. 3**, ROI-level comparisons between illumination conditions were performed using Welch’s *t*-test on *rBFI*, separately for artery and vein ROIs and for each vessel-caliber group (test direction was pre-specified according to the corresponding hypothesis). For multi-group comparisons (*e*.*g*., across vessel-caliber groups), one-way ANOVA followed by Tukey’s post hoc test was applied when appropriate. All statistical analyses were implemented in Python3, primarily using SciPy and stats models. The number of animals, window locations, and ROI/vessel samples used for each analysis is reported alongside the corresponding results/figure legends. No animals were excluded from analysis. Statistical significance was defined as *P* < 0.05. Results are annotated as n.s. (*P* > 0.05), * (*P* < 0.05), ** (*P* < 0.01), *** (*P* < 0.001), and **** (*P* < 0.0001).

## 3. Results

### 3.1 Phototoxicity assessment during conventional prolonged LSCI

To assess phototoxicity-related perfusion changes induced by prolonged standard LSCI illumination, we performed a controlled perfusion imaging experiment after 2 h of standard illumination. As shown in **Fig. 3**, BFI maps exhibit reduced apparent fidelity after prolonged standard illumination, most prominently in small-caliber vessels (<10 μm), with attenuated contrast, blurred vessel boundaries, and a compressed dynamic range in capillaries and terminal arterioles. In the magnified ROIs, several fine vascular branches that are visible under ultra-low illumination become weak or indistinguishable after prolonged standard illumination. This degradation may arise from exposure-associated physiological responses (*e*.*g*., transient vasoconstriction or changes in capillary hematocrit) together with photon-limited measurement noise, as a reduced photon budget lowers the speckle contrast-to-noise ratio. Accordingly, LSCI-derived hemodynamic metrics are jointly influenced by illumination dose, tissue optical properties, and microcirculatory state, and interpretation in photon-limited regimes benefits from consistent calibration and acquisition settings. Notably, the ultra-low illumination dataset in **Fig. 3a** was acquired in a separate session on a different day, with the system re-initialized and the preparation repeated under the same protocol, to avoid carryover from prior irradiation and to ensure that the observed differences primarily reflect the effect of prolonged standard illumination rather than sequential-exposure dependence.

ROI-based quantification in **Fig. 3c-h** further supports these visual observations. After parenchyma normalization, venous ROIs exhibit a pronounced and statistically significant difference in ΔBFI between standard and ultra-low illumination in the small-vessel (<10 μm) class, with a weaker but still detectable difference in the middle-vessel (10-30 μm) class, whereas the parenchymal control and large-vessel (>30 μm) classes show no significant separation (**Fig. 3c**). By contrast, arterial ROIs show smaller or non-significant inter-condition differences across vessel classes (**Fig. 3f**). Vessel stratification by diameter therefore reveals a clear caliber dependence in BFI changes, with the smallest-vessel group showing the largest reduction and larger vessels exhibiting smaller shifts. The class-wise distributions in **Fig. 3d-e** and **Fig. 3g-h** further indicate that this effect is more pronounced in venous than in arterial ROIs. Such bias reduces qualitative interpretability and quantitative comparability across time points or subjects, thereby limiting longitudinal and cross-subject analysis. These findings highlight an urgent need for a computational method that reconstructs physiologically accurate perfusion maps from ultra-low-flux acquisitions, enabling artifact-free, non-invasive, and phototoxicity-safe monitoring of microvascular dynamics.

### 3.2 Optimizing illumination limits of conventional stLASCA

Using the paired acquisition and processing workflow summarized in **Fig. 1**, we evaluate the performance of conventional stLASCA under low exposure. We systematically varied two key parameters: temporal window length (n), i.e., the number of consecutive frames averaged per BFI value; and spatial kernel size (S), i.e., the side length (in pixels) of the square neighborhood for spatial averaging of intensity fluctuations. As shown in **Fig. 4**, this dual-parameter sweep reveals how sensitivity, resolution, and noise robustness trade off under photon-limited conditions. Representative BFI maps (**Fig. 4a-b**) illustrate the classic stLASCA trade-off: increasing S or n improves temporal stability and suppresses speckle noise—yielding smoother perfusion maps—but reduces spatial fidelity. Larger S blurs fine vessels; longer n averages out rapid hemodynamic transients—both degrading microvascular sharpness and capillary–tissue contrast. Under low-exposure, reduced photon flux increases shot noise, which degrades pixel-wise intensity variance estimates. As a result, small vessels—especially those near or below the optical resolution limit—become more prone to artifacts induced by processing parameters. Their visibility now depends not only on anatomical contrast but critically on the trade-off between noise suppression and spatial-temporal resolution. Achieving vascular definition comparable to standard-exposure imaging therefore requires a much higher SNR. This is because stochastic noise obscures fine structures like capillaries and small arterioles. Common compensation strategies—longer exposures, frame averaging, or advanced denoising—introduce trade-offs that can blur or eliminate delicate features such as vessel bifurcations, caliber changes, or microaneurysms.

**Fig. 4.**
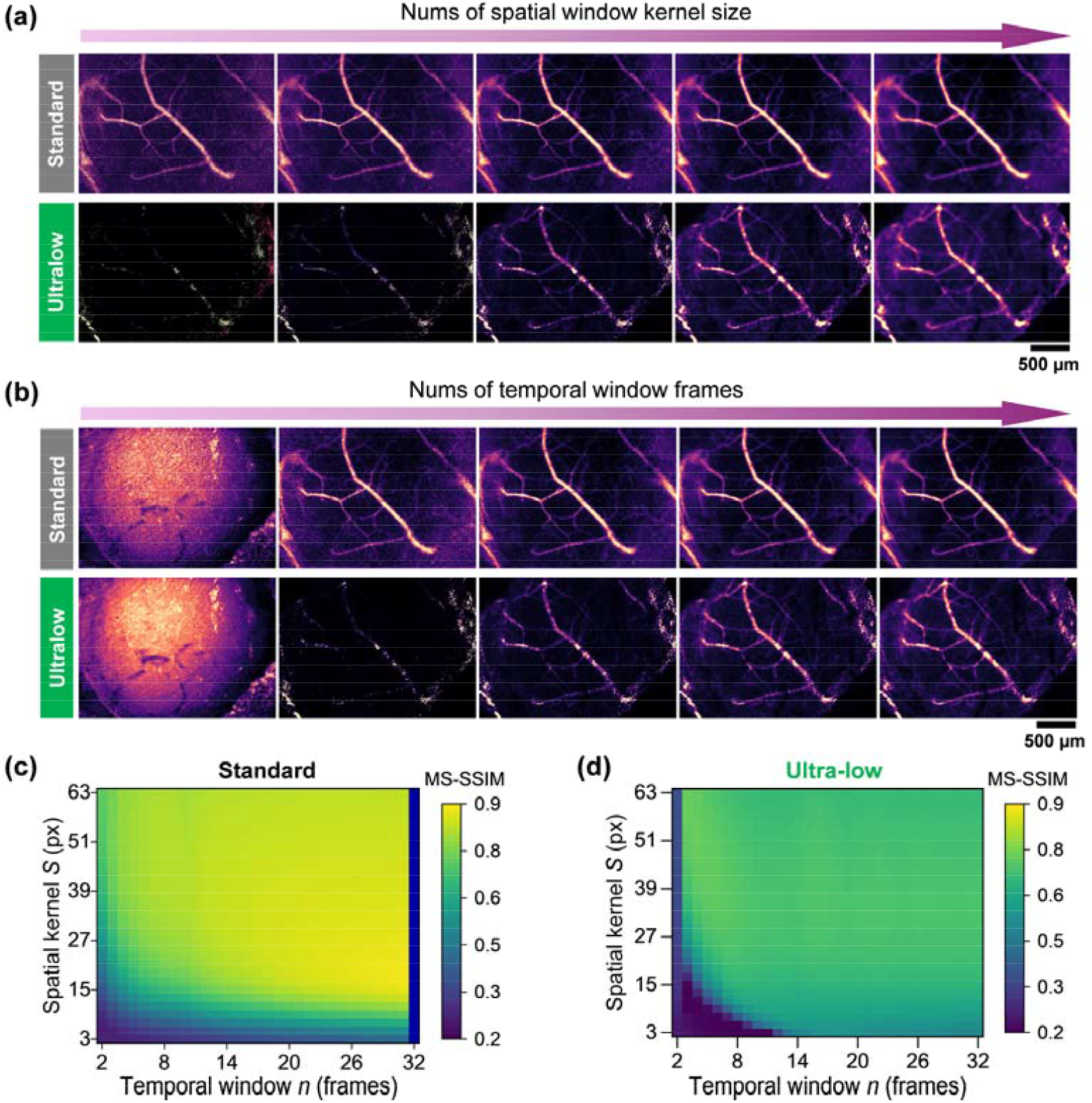
Conventional LSCI algorithm stLASCA falls short under low illumination. (**a**,**b**) BFI maps at ultra-low illumination (1.27 µW/mm^2^, bottom) and standard illumination (∼200 µW/mm^2^, top). Top rows: fixed *n* = 8 with spatial kernel *S* = 3, 7, 15, 31, 63 pixels. Bottom rows: fixed S = 15 with temporal window *n* = 2, 4, 8, 16, 32 frames. Scale bar, 500 µm. (**c**,**d**) MS-SSIM heatmaps versus *n* (2-32) and *S* (3-63), using the standard-illumination reference BFI (*n* = 100, *S* = 15) as baseline.

**Fig. 4c-d** summarizes the dependence of BFI fidelity on temporal window length (*n*) and spatial kernel size (*S*) as 2D MS-SSIM heatmaps. Fidelity is evaluated against the standard-illumination reference BFI (power density = ∼200 µW/mm^2^, *n* = 100, *S* = 15). Under ultra-low illumination (*e*.*g*. power density = 1.27 µW/mm^2^), MS-SSIM is broadly reduced across the (*n*, S) plane, with the largest losses occurring when the estimation is under-averaged (small n and/or small *S*) and thus dominated by photon shot noise. Increasing n improves stability but reduces temporal responsiveness, whereas increasing S suppresses speckle noise at the cost of attenuating fine microvascular structures; consequently, the heatmaps exhibit an operating region where fidelity is maximized rather than a monotonic improvement with smoothing. Overall, the parameter-dependent patterns indicate that photon-limited measurements retain structured vascular information, but conventional stLASCA recovers it only through averaging that inevitably trades spatial/temporal resolution. These observations motivate a learning-based reconstruction that leverages spatiotemporal priors to suppress shot-noise–dominated fluctuations while preserving vessel-level morphology and contrast under extreme photon constraints.

### 3.3 TunLSCI: Few-frame TransUNet-based recovery for ultra-low-illumination LSCI

We evaluated whether TunLSCI can reconstruct high-fidelity BFI maps from ultra-low-illumination speckle sequences with minimal temporal support, achieving accurate perfusion recovery from as little as a single frame (*n* = 1-16), as required under stringent frame-budget conditions imposed by residual motion and the need for high temporal resolution in awake imaging. Using *in vivo* mouse CBF imaging data (**Fig. 5**), we benchmarked performance across frame counts (*n* = 1, 4, 8, 16), all acquired at identical fluence (≈0.17 µJ/mm^2^/frame) and processed with fixed stLASCA parameters (spatial kernel size S = 15). Conventional stLASCA reconstructions (**Fig. 5a**) show smoother, more stable BFI estimates as n increases—consistent with ensemble averaging—but even at *n* = 16, noise persists in low-flow regions, and systematic bias remains: high-flow vessels are attenuated, while low-flow parenchyma shows inflated BFI due to the nonlinearity and outlier sensitivity of traditional speckle contrast computation.

**Fig. 5.**
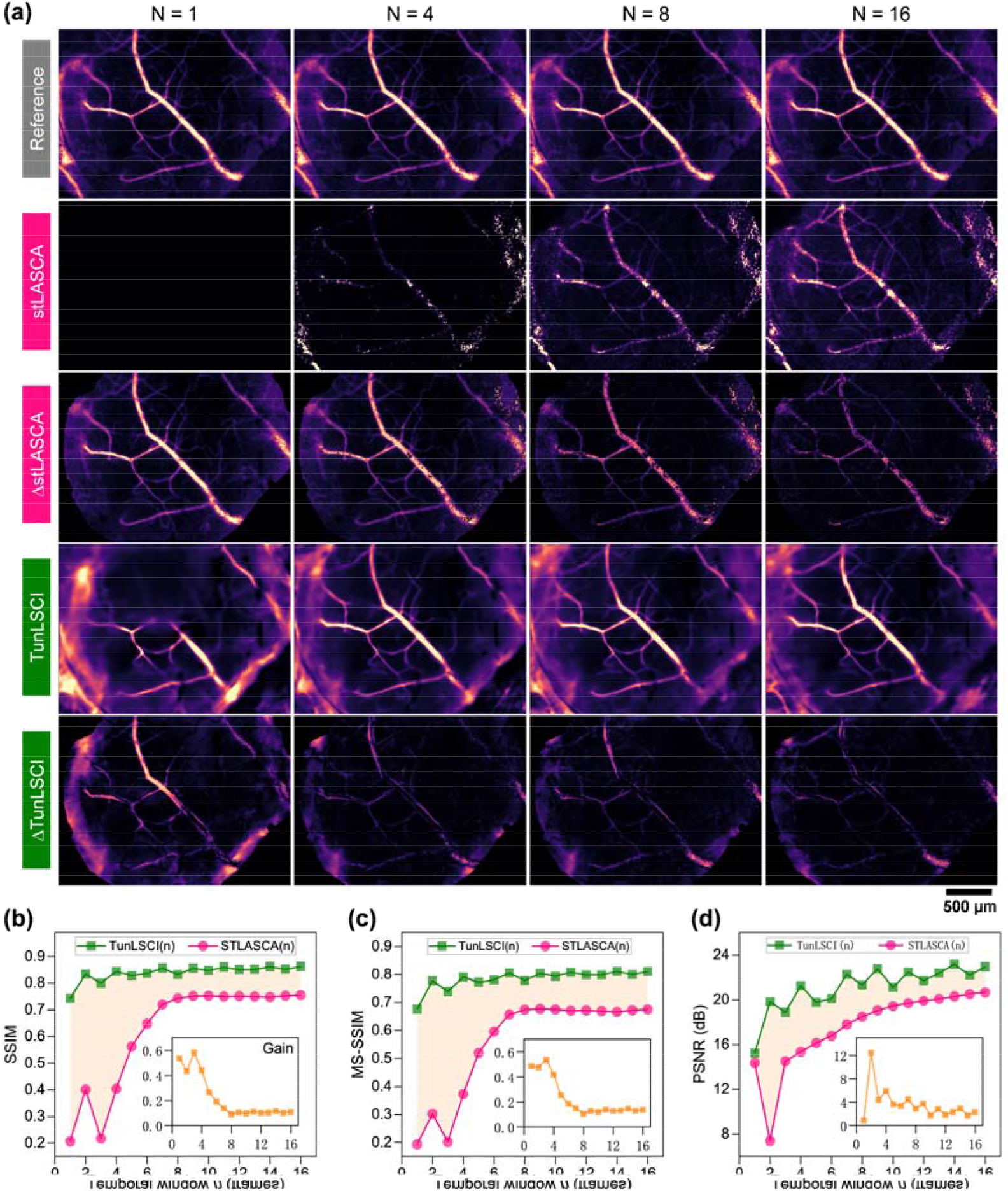
Few-frame recovery performance under low-exposure acquisition. (**a**) Representative maps from one FOV: reference BFI (stLASCA, power density = 200 µW/mm^2^, *n* = 100, *S* = 15), low-exposure stLASCA(*n*) (power density = 1.27 µW/mm^2^, n = 1/4/8/16, S = 15), TunLSCI(n) predictions, and absolute error |*I* _*pred*_ - *I*_*Ref*_|. Scale bar, 500 μm. (**b-d**) SSIM, MS-SSIM, and PSNR versus *n* = 1-16 on 8-bit BFI within the vessel mask for low-exposure stLASCA(n) and TunLSCI(n).

In contrast, TunLSCI(*n*) maintains vascular continuity and resolves fine capillary branching accurately across all photon-counting levels (*n* = 1-16), while robustly suppressing multiplicative speckle noise—a key challenge in low-light laser speckle contrast imaging. Unlike conventional stLASCA, which blurs microvessels or creates artificial discontinuities under low-photon conditions, TunLSCI preserves anatomical fidelity without compromising noise resilience. Its absolute-error maps—computed as pixel-wise differences between predicted and ground-truth perfusion maps—show residuals confined to vessel edges or intensity transitions; no global warping, topological errors, or large-scale distortions occur (**Fig. 3a**). Quantitatively, TunLSCI outperforms conventional stLASCA in SSIM, MS-SSIM, and PSNR across all *n* = 1-16 (**Fig. 3b-d**). Gains are largest at the lowest n (1-4), where stLASCA suffers from high variance and over-smoothing—degrading dynamic contrast, flow sensitivity, and boundary sharpness. At *n* =3, TunLSCI improves SSIM by >0.5 and PSNR by >5 dB versus stLASCA, demonstrating superior signal recovery from near-noise-floor data.

### 3.4 TunLSCI is more robust than stLASCA under low-exposure conditions

Ultra-low-illumination LSCI acquisitions frequently require exposure-time adjustments to accommodate session-to-session variations in speckle brightness, tissue reflectance, and camera dynamic range, even when the illumination power density is held constant. To verify exposure-time robustness, the TunLSCI model was trained using only 60-ms low-power inputs (power density = 1.27 µW/mm^2^) and evaluated on two independent test conditions acquired in the same FOV: unseen 60-ms recordings and 120-ms recordings (2× exposure). The reference target definition was kept fixed as the standard-illumination stLASCA BFI (power density ∼200 µW/mm^2^, *n* = 100, S = 15), ensuring that performance differences reflect exposure-time variation rather than any change in target construction. As shown in **Fig. 6a**, at representative frame budgets (*n* = 4 and *n* = 8), TunLSCI preserves vessel continuity and microvascular morphology under both exposure settings, whereas stLASCA exhibits more pronounced exposure-dependent changes in background texture and small-vessel appearance. Quantitatively, **Fig. 6b-d** reports SSIM, MS-SSIM, and PSNR within the vessel mask across *n* =1-16 frames, where TunLSCI consistently achieves higher fidelity than stLASCA for both 60-ms and 120-ms inputs, with the most substantial gains at the most photon-limited frame budgets (*e*.*g*., at *n* = 1: MS-SSIM/PSNR = 0.705/16.64 dB vs 0.192/14.35 dB (60 ms); 0.715/16.58 dB vs 0.192/14.35 dB (120 ms)). Importantly, **Fig. 6e-g** directly quantifies exposure-time sensitivity by plotting the metric differences between 120 ms and 60 ms. In this exposure-gap analysis, TunLSCI exhibits only minor metric drift (<0.04 SSIM variation and <2.3 dB PSNR shift), whereas conventional stLASCA shows substantially larger degradation (>0.3 SSIM drop and >3.2 dB PSNR loss). Together, these results support stable vessel-level BFI recovery across exposure-time variations under ultra-low illumination, without relying on aggressive spatiotemporal averaging.

**Fig. 6.**
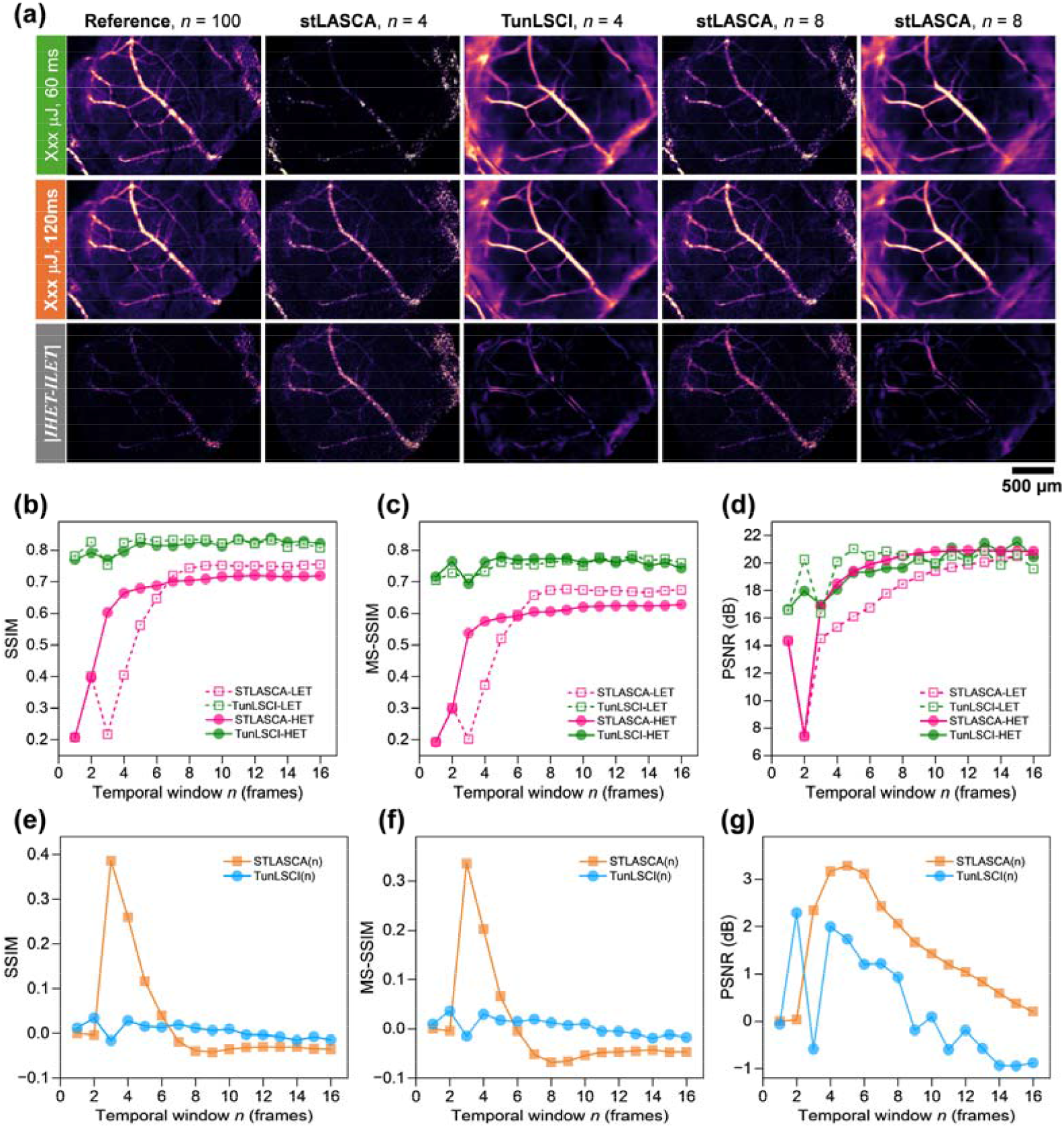
Robustness to ultra-low illumination exposure-time variation (60 ms vs 120 ms). (**a**) Representative maps at *n* = 4 and *n* = 8: stLASCA(*n*), TunLSCI(*n*), reference, and long–short exposure difference maps |*I*_120 ms_ - *I*_60 ms_|. Metrics were computed within the vessel mask on 8-bit BFI maps. Scale bar, 500 µm. (**b-d**) SSIM, MS-SSIM, and PSNR versus *n* = 1-16 for stLASCA(*n*) and TunLSCI (*n*) under both exposure conditions. (**e-g**) Exposure-gap curves for each metric. TunLSCI trained on 60-ms low-exposure data and evaluated on 60-ms and 120-ms low-power recordings. Reference BFI: standard-exposure stLASCA (*n* = 100, S = 15) from the paired recording. TunLSCI outputs were visualized with a unified color scale.

## 4. Discussion

Integrating deep learning into LSCI alleviates the long-standing trade-off between image quality and illumination dose in prolonged monitoring by enabling informative reconstructions from photon-limited measurements [23, 24]. Unlike traditional LSCI—which relies on high laser exposure to ensure a sufficient SNR—our TunLSCI reduces phototoxicity and tissue damage [25] and cuts laser power density by ∼100× versus prior methods (**Fig. 7**). Our method combine few-frame denoising, structure-preserving restoration and physics-aware inference for time-/multi-exposure and system-specific acquisitions, collectively improving blood-flow readouts under photon-limited and heterogeneous imaging conditions. It minimizes photon flux without sacrificing spatial or temporal resolution, enabling longer-term neurovascular and microcirculatory studies. Using hierarchical feature learning, it suppresses photon-limited noise while preserving hemodynamically critical information. It can also reduce motion artifacts—especially vital in awake animal studies—by enforcing temporal coherence in the loss function. In addition, exposure-aware learning modules implemented as software-layer post-processing can be integrated into existing pipelines to enable near real-time flow inference across exposure settings [26-28].

**Fig. 7.**
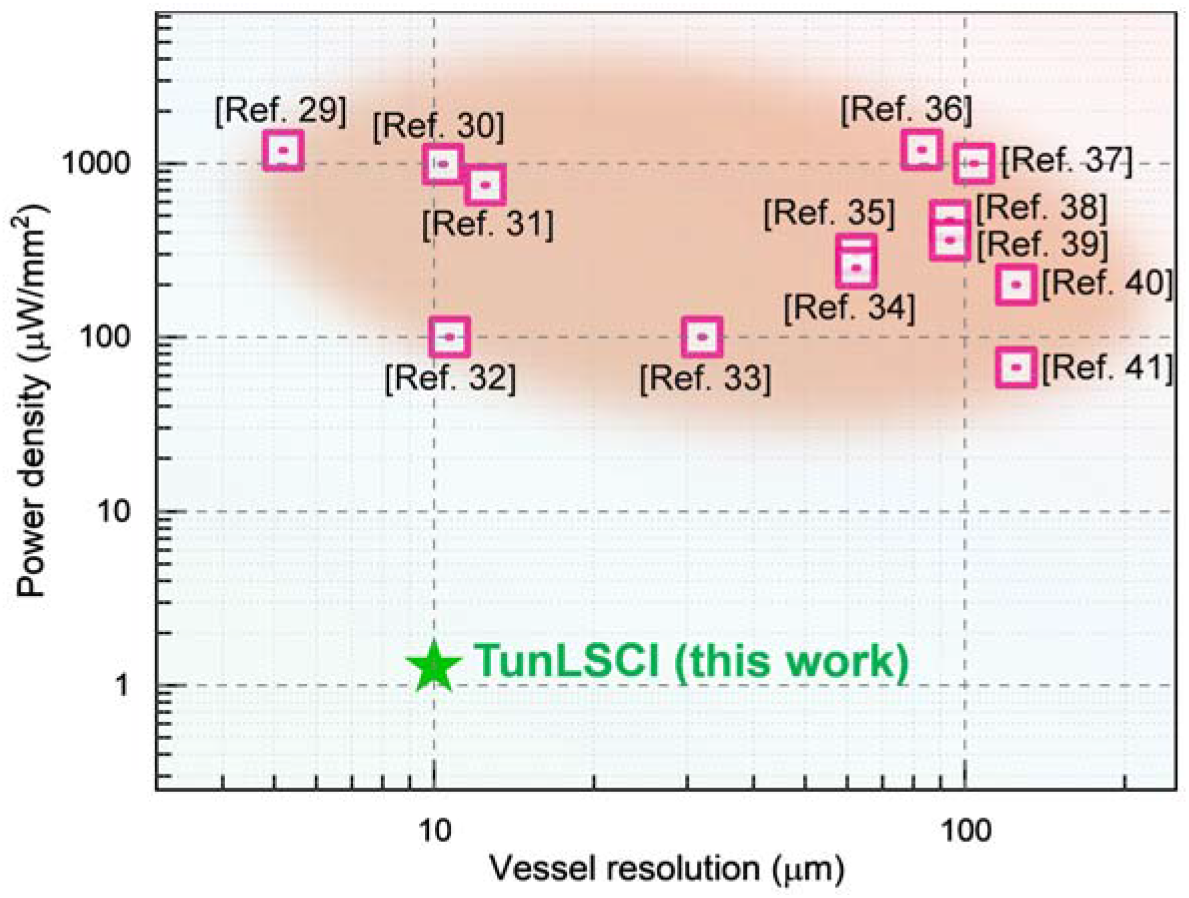
Comparison of the vessel image resolution and laser illumination power density of TunLSCI with those of previously reported state-of-the-art LSCI measurements reported in the past 10 years. All data points were obtained from the studies presented in the aforementioned descriptions and additionally listed in [29-41].

Despite these advances, significant limitations persist. For instance, the cross-species generalizability of TunLSCI is yet to be validated. For example, it’s reported that the unsupervised methods can introduce spectral biases in speckle contrast enhancement, distorting hemoglobin oxygen saturation estimates [42]. The interplay between laser wavelength and DL enhancement is needed to be further characterized — networks trained at 785 nm may underperform at 850 nm due to differences in scattering regimes [35]. And ethical considerations surrounding computational phototoxicity—whereby aggressive denoising inadvertently masks genuine tissue damage—have yet to be addressed through large-scale controlled trials. Future systems could couple speckle data with auxiliary biomarkers (*e*.*g*., NADH autofluorescence) to ground-truth DL enhancements against metabolic states [43]. Additionally, self-supervised frameworks that continuously recalibrate based on real-time tissue feedback would address domain shift issues while minimizing expert annotation burdens [44]. From a hardware perspective, the emergence of single-photon avalanche diode (SPAD) arrays with time-gating capabilities will enable diffusion models to resolve depth-specific flow dynamics at unprecedented scales [26]. On the algorithmic front, explainable AI methodologies like attention mapping could demystify DL decisions, fostering clinician trust [45].

As these methods mature through interdisciplinary collaboration, they will advance our understanding of neurovascular coupling, microcirculatory dysfunction, and intraoperative hemodynamics—safely and without iatrogenic risk. By prioritizing biological fidelity alongside technical precision, DL-enhanced LSCI remains a trustworthy tool for exploring brain function, microvascular diseases, and therapeutic interventions.

## Acknowledgment

The Institutional Animal Care and Use Committee (IACUC) of Shanghai Institute of Optics and Fine Mechanics (No.2025-001) approved the animal experiment protocol.

## Data availability

Data supporting the findings are available from the corresponding authors upon reasonable request.

